# Persistent activity during working memory maintenance predicts long-term memory formation in the human hippocampus

**DOI:** 10.1101/2024.07.15.603630

**Authors:** Jonathan Daume, Jan Kamiński, Yousef Salimpour, William S. Anderson, Taufik A. Valiante, Adam N. Mamelak, Ueli Rutishauser

## Abstract

Working Memory (WM) and Long-Term Memory (LTM) are often viewed as separate cognitive systems. Little is known about how these systems interact when forming memories. We recorded single neurons in the human medial temporal lobe while patients maintained novel items in WM and a subsequent recognition memory test for the same items. In the hippocampus but not the amygdala, the level of WM content-selective persist activity during WM maintenance was predictive of whether the item was later recognized with high confidence or forgotten. In contrast, visually evoked activity in the same cells was not predictive of LTM formation. During LTM retrieval, memory-selective neurons responded more strongly to familiar stimuli for which persistent activity was high while they were maintained in WM. Our study suggests that hippocampal persistent activity of the same cell supports both WM maintenance and LTM encoding, thereby revealing a common single-neuron component of these two memory systems.

## Introduction

Working Memory (WM) is the ability to hold and manipulate a small amount of information “in mind”, an ability that is fundamental to many aspects of cognition (Baddeley 2012). Since at least the 1960s, when Atkinson and Shiffrin’s first proposed their model of memory (Atkinson and Shiffrin 1968), it has been theorized that WM (then called Short-Term Memory) and Long-Term Memory are two separated but connected systems. This model and later theories of WM suggest that WM acts as intermediary between perception and LTM (Baddeley 2003), a relationship that has been studied extensively for decades. Indeed, in many instances, information held in WM is encoded better into LTM compared to information not held in WM. Behaviorally, this relationship has been shown in many studies (Hartshorne and Makovski 2019). For instance, words which are maintained longer in WM were later recalled better (Souza and Oberauer 2017), and items stored in WM were better remembered in a surprise recognition test compared to items only attended or passively viewed (Daume et al. 2017). A recent meta-analysis and new experiments show that the impact of holding information in WM on the quality of LTM is especially strong in the visual domain (Hartshorne and Makovski 2019).

Despite the ubiquity of WM-LTM interactions seen behaviorally, little is known about the mechanisms by which the two memory systems interact. fMRI (Davachi et al. 2001; Schon et al. 2004; Ranganath et al. 2005; Blumenfeld and Ranganath 2006; Axmacher et al. 2008), scalp M/EEG (Khader et al. 2007, 2010; Daume et al. 2017), and intracranial EEG studies (Axmacher et al. 2009, 2010b) indicate that the extent of activation of a given part of the brain as assessed by BOLD-fMRI or oscillatory power during WM maintenance can be predictive of both WM maintenance success and LTM encoding success. Further, in dual task paradigms (Axmacher et al. 2009, 2010b), high WM demands disrupt LTM encoding processes, arguing that they are not independent. Overall, these findings indicate that the neuronal substrate of the two processes are at least partially overlapping or shared, and are located in the same areas of the brain. However, it remains unclear what exactly is shared in terms of the neuronal substrate. One possibility, motivated by theoretical models (see below), is that the two processes of WM encoding and LTM encoding engage the same cells, but this prediction has not been tested experimentally so far.

A hypothesis that has motivated a large body of work is that the sustained maintenance of memoranda in WM enables the gradual strengthening of LTM traces through synaptic plasticity (Hebb, 1949). A key prediction from this model is that the stronger the activation of the neurons that represent memory content during WM maintenance, the stronger the resulting LTM. Here we test this hypothesis directly by recording from WM memory-content selective neurons and assess their relationship to LTM encoding in a task with trial-unique novel stimuli for which we later test memory strength. We note that this design is different from most WM studies, in which the items held in WM are re-used throughout an experiment.

Neuroimaging, iEEG, and behavioral studies indicate that the medial temporal lobe, and in particular the hippocampus, are strong candidates for shared WM-LTM processes (Squire 2004). Indeed, recent studies indicate that the MTL, particularly the hippocampus, is critical for both WM and LTM in many circumstances as assessed by behavior and neural activity (Nichols et al. 2006; Piekema et al. 2006; Axmacher et al. 2010a; Jeneson and Squire 2012; Libby et al. 2014; Leszczyński et al. 2015). The MTL has therefore emerged as a key candidate for the area where the WM and LTM system might interact.

Transient maintenance can only strengthen LTM through synaptic plasticity as proposed by Hebb (Hebb 1949) if the neurons involved in WM maintenance are also part of the circuit that encodes LTM. A candidate substrate that fulfills these criteria is memoranda-selective persistently active neurons. Such cells, which constitute a relatively well understood cellular substrate for maintaining information in WM, have been documented for highly familiar stimuli (Funahashi et al. 1989; Chafee and Goldman-Rakic 1998; Rainer et al. 1998; Kamiński et al. 2017; Kornblith et al. 2017; Daume et al. 2024). In humans, WM-content selective persistently active neurons have been described in the MTL. The activity of these cells is behaviorally relevant and scales with memory load during maintenance of their preferred stimuli (Kamiński et al. 2017; Kornblith et al. 2017; Boran et al. 2019; Daume et al. 2024). We hypothesize that these cells might contribute to both WM maintenance and LTM encoding, thereby allowing the transient maintenance of activity to translate into structural changes through synaptic plasticity. If so, the extent of persistent activity should be indicative of later LTM memory strength as assessed behaviorally and/or neuronally. Here, we used the opportunity to invasively record single neurons in the human MTL in patients undergoing invasive epilepsy monitoring with depth electrodes to test this hypothesis. Patients performed both a WM and an LTM task in the same recording sessions, with a shared stimulus set between the two tasks. This design allowed us to assess whether maintaining a given trial-unique item in WM influences how well that item will later be remembered in a recognition memory test.

## Results

41 patients (48 sessions; 1 non-binary; 20 females; 20 males) performed a modified Sternberg WM task with novel images, followed by a subsequent LTM recognition test. In the WM task, patients were asked to hold either one (load 1) or three (load 3) sequentially presented images in their mind until a probe picture appeared 2.5 – 2.8 s later (**Fig. 1a top**) (Daume et al. 2024). The task was to indicate whether the probe picture was identical to one of the encoding pictures just presented before the delay period or not. All pictures shown during encoding were novel, i.e. never shown to the subject before. The images shown during the probe were always familiar to the subject, either from having seen them in the current trial during encoding or during a previous trial (an image is shown twice at most, if used as probe; see Methods). Images are drawn from five different picture categories (people, animals, cars (or tools depending on version), food, landscapes). After a delay of 10-30 min, patients performed a LTM recognition task in which half of the images shown were the same as those shown during the WM task (familiar items) and half were novel (**Fig. 1a bottom**). In this part of the task, subjects indicated whether a given picture was “old” (i.e., seen in the earlier performed WM task) or “new”. Patients were also asked to indicate the confidence in their response, i.e., how confident they were that a given item was old or new (“sure”, “unsure”, “guessing”).

**Figure 1.**
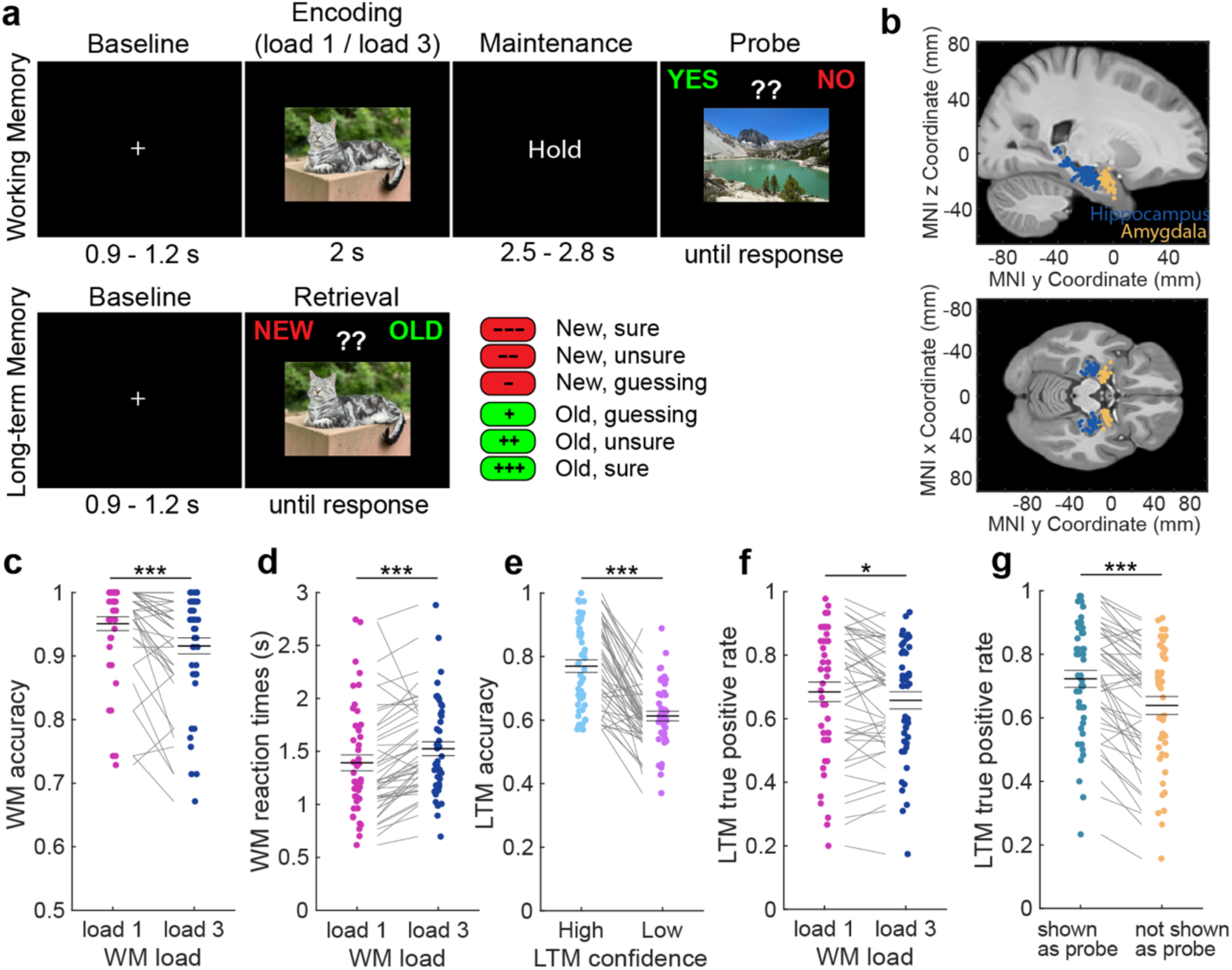
Experimental design and behavioral analysis. **(a)** The study contained a WM part (top) and a subsequent LTM recognition part (bottom). In the WM task, patients had to encode either one (load 1) or three (load 3) pictures in their WM and to maintain these items until a probe picture appeared a few seconds later. Their task was to indicate whether the probe was part of the encoded items in a given trial or not. All encoding pictures were novel and drawn from five different picture categories. In the LTM task, occurring after a 10-30 min break, patients had to answer whether each presented item on the screen was old (i.e., seen in the previous WM task) or new while indicating their confidence in their response. **(b)** We recorded single neuron activity from the hippocampus and the amygdala of 41 patients across 48 sessions. Each dot is a patient. **(c,d)** WM behavior. Patients performed **(c)** more accurate and **(d)** faster in load 1 compared to load 3 trials. **(e-g)** LTM behavior. **(e)** LTM was more accurate in high than low confidence trials. **(f)** Items previously maintained in load 1 trials were remembered more accurately than when maintained in load 3 trials. **(g)** Pictures that were used as probe and therefore presented twice were remembered better than items not shown as probe. In **(c-g)** we used permutation-based t-tests. Center lines represent mean ± s.e.m. Each dot is a session. * p < 0.05; *** p < 0.001.

Patients performed well in both parts of the task, with an average accuracy of 93.3 ± 7.6 % (mean ± SD) in the WM task and 70.3 ± 9.8 % in the LTM task, respectively (both p < 0.0001 as compared to 50% chance; permutation-based t-test; 2 sessions from two different patients were excluded from all analyses due to an accuracy of less than 55% in at least one of the tasks; see **Table S1;** see **Fig. S1** for behavior results of 100 healthy participants). In the WM task, patients performed with higher accuracy (**Fig. 1c**; load 1 – load 3: t(45) = 4.89, p < 0.0001; permutation-based paired t-tests were used throughout the manuscript unless stated otherwise; t-values are provided as reference only; see Methods) and faster (**Fig. 1d**; t(45) = −4.29, p < 0.0001) in load 1 than load 3 trials. In the LTM task, retrieval was more accurate (**Fig. 1e**; t(44) = 8.66, p < 0.0001; one patient did not use confidence ratings and was therefore excluded from all confidence-related analyses) and faster (mean RT high: 1.83 ± 0.57 s; mean RT low: 2.62 ± 0.66 s; t(44) = - 11.13, p < 0.0001) when patients indicated high confidence in their responses. Low confidence was defined as an average of “unsure” and “guessing” responses. Items that were encoded in load 1 trials in the previous WM task were remembered better than items encoded during load 3 trials (**Fig. 1f**; t(45) = 2.42; p = 0.018). Moreover, items that were also used as the probe and therefore presented twice were remembered better than items not used as the probe (**Fig. 1g**; t(45) = 7.25, p < 0.0001).

We recorded single neuron activity from the hippocampus and the amygdala while patients performed the two tasks (**Fig. 1b**). In total, 883 single units across both brain areas were included in our analyses, 351 from the hippocampus and 532 from the amygdala (see Methods). The same units were recorded for both tasks. The analysis presented here is partly based on a previously published dataset (Daume et al. 2024), but in that report only the WM task was analyzed. Further, more subjects were added in the present manuscript. The LTM part that is the focus of this paper is unpublished. Spike sorting results were assessed quantitatively (**Fig. S2**). We use the terms neuron, unit, and cell interchangeably to refer to a putative single neuron.

### Category neurons remain persistently active when their preferred category is held in WM

We first selected for neurons whose response following stimulus onset differed significantly between the category of the images shown. We refer to such neurons as ‘category neurons’ throughout (we note that this type of cell is also referred to as visually selective (VS) in other papers (Rutishauser et al. 2015, 2021; Bausch et al. 2021); The two terms are equivalent for purpose of this study). To select category neurons, we assessed whether the firing rate (FR) in a 200 – 1200 ms window following picture onset (encoding 1-3 & probe) was significantly correlated with the five possible picture categories (1×5 ANOVA, followed by a right-sided permutation-based t-test between the category with maximal average spike count and all other categories; if both tests were p < 0.05, we classified a neuron as a category neuron with the preferred category being the one with maximal average spike count; see Methods). As shown previously, category neurons remain persistently active during the maintenance period of the WM task when their preferred picture is held in WM (Daume et al. 2024). Selecting for category neurons during picture presentation leaves their FRs during the maintenance period of the task independent for subsequent statistical analyses. In the hippocampus, 104 (29.65 %) neurons qualified as category neurons, and in the amygdala 220 (41.35 %) neurons qualified as category neurons (see **Fig. 2a,d** for example neurons from each area).

**Figure 2.**
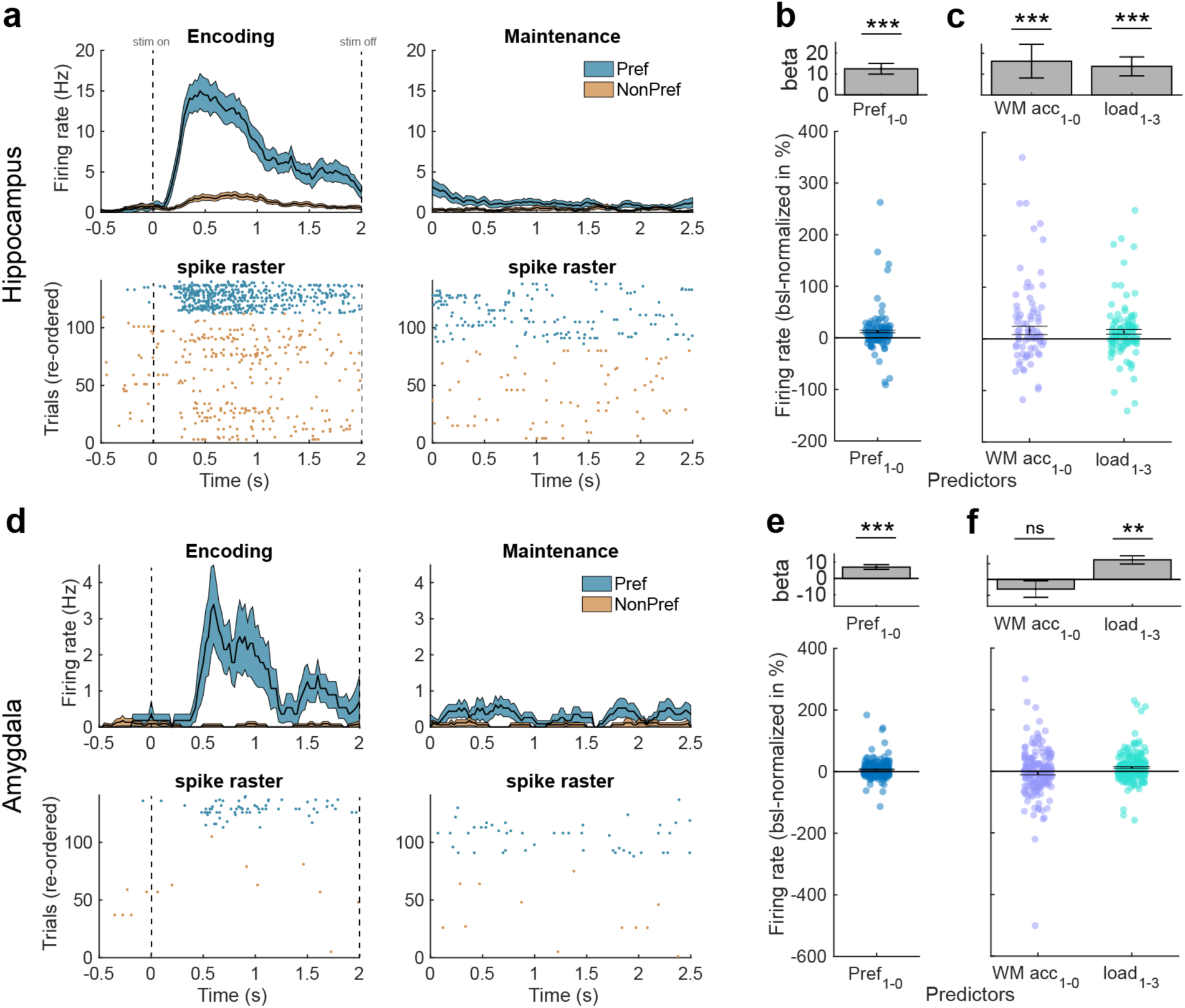
Category neurons. **(a-c)** Characterization of category neurons in the hippocampus. **(a)** Example category neuron. The preferred category of this neuron was “animals”. Top: Peri-stimulus time histogram (PSTH, bin size = 200 ms, step size = 25 ms) during the first picture in each correct trial. Colored areas represent ± s.e.m. Bottom: raster plot with trials re-ordered into preferred and non-preferred categories. Stimulus and maintenance onset is at t = 0s (left and right, respectively). **(b)** Firing rates of category neurons from the hippocampus remained persistently active and were higher for preferred than unpreferred trials during the WM maintenance period. Top: Beta value extracted from the GLM for preferred/non-preferred regressor in units of “percent change to baseline” (−900 – −300 ms before first picture onset). Bottom: Distribution of FR differences between preferred (pref = 1) and non-preferred (pref = 0) trials across all hippocampal category neurons (each dot is a neuron, n = 104). FRs were baseline-normalized to represent percent change to baseline. **(c)** When the preferred category remained in WM, category neurons in the hippocampus had higher FRs in correct as compared to incorrect and in load 1 as compared to load 3 trials. **(d-f)** Characterization of category neurons in the amygdala. **(d)** Same as in (a) but for an example category neuron from the amygdala. **(e)** Firing rates of category neurons from the amygdala also remained persistently active and were higher for preferred than unpreferred trials during the WM maintenance period (n = 220). **(f)** Their FRs were higher in load 1 as compared to load 3 trials when their preferred category was maintained but there was no difference between correct and incorrect trials. In **(b,c,e,f)** we computed mixed-effects GLMs. Error bars represent standard errors of the coefficient. In **(b,e)** each dot is a neuron. ** p < 0.01, *** p < 0.001, ns = not significant.

To confirm that category neurons remained persistently active during the maintenance period, we computed a mixed-effects generalized linear model (GLM) using *preferred category* (2 levels, true/untrue, categorical) as fixed effect and *neuron ID* nested into *patient ID* as random intercept in each area (these random intercept terms were used for all GLMs; see Methods). We used baseline-normalized FRs from all correct WM trials during the maintenance period (0-2.5 s after last encoding picture offset) for this analysis. In the hippocampus, category neurons remained persistently active throughout the maintenance period (intercept: beta = 17.86, p = 1.38 × 10^−3^; mixed-effects GLM) and had significantly higher FRs during trials in which images from the preferred category were maintained in WM (**Fig. 2b**; preferred category: beta = 12.47, p = 8.82 × 10^−7^). Using only preferred trials and modelling WM *accuracy* (2 levels, WM correct/incorrect, categorical) as well as *load* (2 levels, load 1/load 3, categorical) as fixed effects, we further observed that category neurons in the hippocampus had higher FRs in (1) correct than incorrect trials (beta = 16.22, p = 0.046) and (2) in load 1 than in load 3 trials (**Fig. 2c**; beta = 13.769, p = 0.0023).

In the amygdala, category neurons also remained persistently active throughout the maintenance period across all correct trials (intercept: beta = 9.78, p = 6.76 × 10^−4^) with FRs higher for preferred than unpreferred categories (**Fig. 2e**; preferred category: beta = 6.91, p = 9.38 × 10^−7^). However, unlike in the hippocampus, using only preferred trials category neurons in the amygdala showed a significant main effect only for *load* (**Fig. 2f**; load 1 – load 3; beta = 12.309; p = 2.46 × 10^−7^), but the effect for WM *accuracy* (correct – incorrect: beta = −6.13; p = 0.24) was not significant.

### Persistent activity of category neurons in the hippocampus - not the amygdala - predicts LTM formation

We next sought to investigate whether the activity of category neurons during the maintenance period of the WM task predicted the success of LTM formation. To that end, we computed a mixed-effects GLM with baseline-normalized FRs from the maintenance period. For a given neuron, we used the subset of trials for which the subject answered the WM question correctly and for which images of the preferred category were tested in the LTM recognition task (not all images seen during WM were shown during the LTM test). We modelled *LTM accuracy* of each image (2 levels, remembered/forgotten, categorical), *confidence* (3 levels, sure/unsure/guessing, continuous), *brain area* (2 levels, hippocampus/amygdala, categorical), as well as their interactions as fixed effects. We did not observe any significant main effects nor was the interaction between *LTM accuracy* and *confidence* significant (**Fig. 3a**; all p > 0.26). However, all interaction terms including *area* showed significant modulations (confidence x area: beta = 16.88, p = 0.025; LTM accuracy x area: beta = 59.78, p = 0.0030; confidence x LTM accuracy x area: beta = −28.64, p = 0.0029), suggesting that the relationship between FR and *LTM accuracy* and *confidence* differed between the hippocampus and the amygdala. We therefore repeated the analysis in each area separately. This revealed a significant main effect of *confidence* (**Fig. 3b**; beta = 20.30, p = 4.17 × 10^−3^) and *LTM accuracy* (beta = 67.90, p = 4.43 × 10^−4^), as well as a significant interaction between the two terms (beta = −30.80, p = 7.74 × 10^−4^) in the hippocampus but not in the amygdala (confidence: beta = 1.54, p = 0.69; LTM accuracy: beta = −2.27, p = 0.82; confidence x LTM accuracy: beta = 1.02, p = 0.83). These results suggest that activity during the maintenance period of the WM task of category neurons from the hippocampus were higher for later remembered than forgotten trials and therefore predictive of later LTM retrieval performance. This was not the case for persistent WM activity of category neurons from the amygdala.

**Figure 3.**
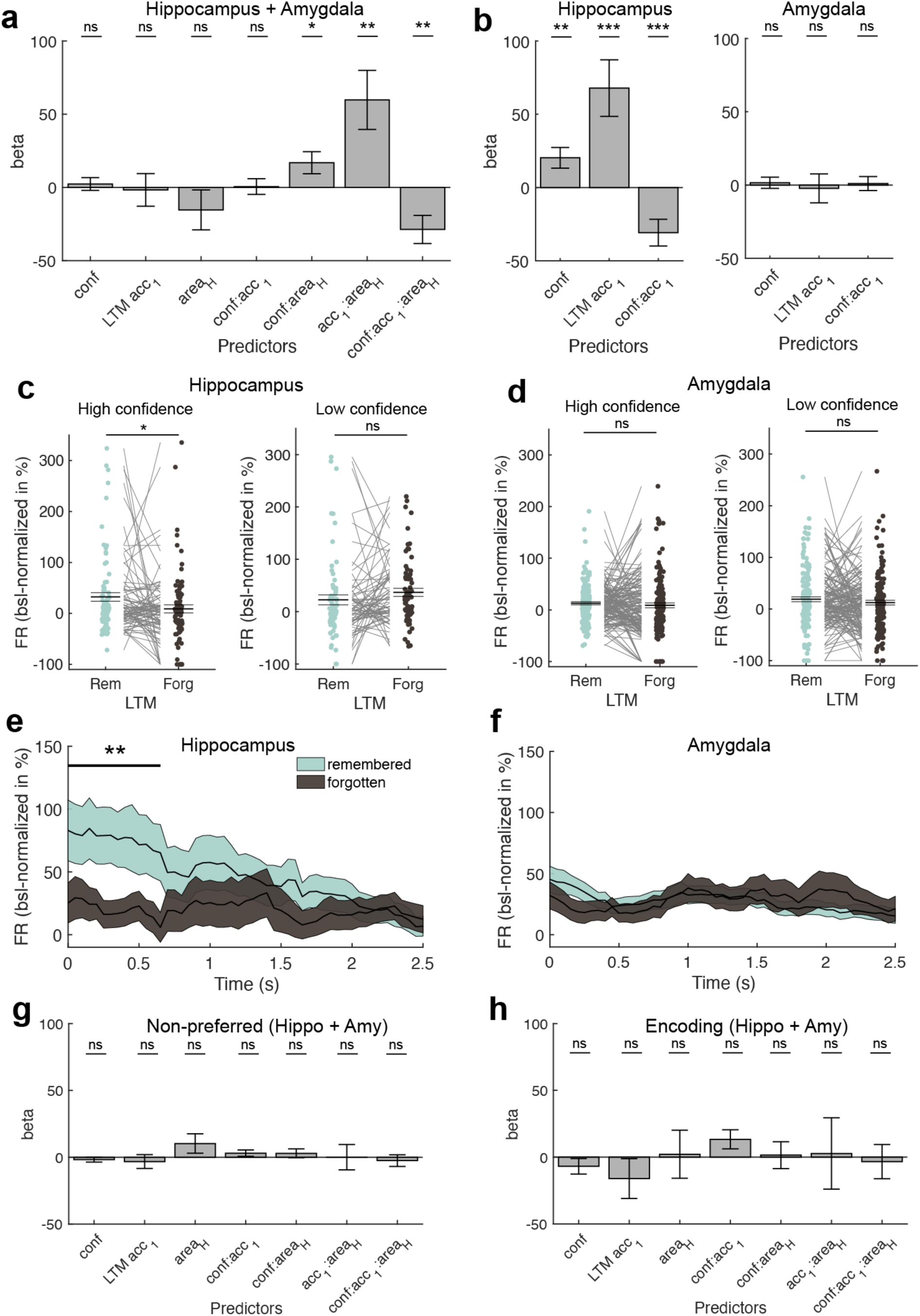
Relationship between WM maintenance activity and LTM formation. **(a)** Mixed-effects model using the FR obtained during the WM maintenance period in correct and preferred trials of all category cells across both regions, modelling *confidence,* subsequent *LTM accuracy (*remembered vs forgotten*), area*, and their interactions as fixed effects and *neuron ID* nested into *patient ID* as random intercepts. We found significant modulations of FR by interactions of *confidence* and *LTM accuracy* with *area*, suggesting differences in FR modulations by LTM accuracy and confidence per area. **(b)** Mixed-effects GLM results separately for the hippocampus (left) and the amygdala (right). Only in the hippocampus, we observed that persistent activity during the WM maintenance period predicted later LTM accuracy as well as confidence. **(c,d)** Comparison of FRs during the maintenance period between later remembered and forgotten images, separately for high (left) and low confidence trials (right) for category neurons from **(c)** the hippocampus and **(d)** the amygdala. Statistics are permutation-based t-tests, each dot is a neuron. FRs differed significantly between remembered and forgotten images in the hippocampus for high but not low confidence trials. **(e,f)** Time-resolved FR differences between later high-confident remembered and forgotten trials in (e) the hippocampus and (f) the amygdala for the maintenance period. FR differences between remembered and forgotten trials in the hippocampus were strongest in the first section of the WM delay period (0 – 650 ms). Cluster-based permutation t-test. Colored areas represent ± s.e.m. t=0 marks the onset of the maintenance period. **(g)** Mixed-effects model results using the WM maintenance period FRs of non-preferred trials across all category neurons from both regions. **(h)** Mixed-effects model results using the FRs during the first picture presentation (encoding 1; 0-2 s after picture onset; preferred images and correct trials only) across all category neurons from both regions. In **(a,b,g,h)** error bars represent standard errors of the coefficients. Betas are shown in units of baseline-normalized FRs (percent change to baseline, −900 – −300 ms before first picture onset). In **(c,d)** center lines represent mean ± s.e.m. * p < 0.05, ** p < 0.01, *** p < 0.001, ns = not significant.

In each area, we next tested FRs across all category neurons from the maintenance period of the WM task between later remembered and forgotten separately for high and low LTM retrieval confidence. In the hippocampus, FRs were higher for remembered than forgotten trials for high confidence (**Fig. 3c**; t(82) = 2.16, p = 0.028; some neurons were removed due to insufficient data in at least one of the conditions or since they differed ±3 SD from the mean across all neurons and conditions; see Methods), but not low confidence trials (t(68) = −1.38, p = 0.17). In the amygdala, we neither observed a significant effect for high (**Fig. 3d**; t(158) = 0.84, p = 0.41) nor low confidence trials (t(152) = 1.15, p = 0.26).

To observe when during the maintenance period of the WM task the effect between later remembered and forgotten trials was present, we tested time-resolved FRs between remembered and forgotten trials for all high-confident trials separately for category neurons from hippocampus and amygdala. In the hippocampus, FRs during the beginning of the maintenance period (0 – 650 ms) differed significantly between the two conditions (**Fig. 3e**; cluster-p = 0.0042; cluster-based permutation test). In the amygdala, we did not observe any significant cluster throughout the entire maintenance period of the WM task (**Fig. 3f**; cluster-p = 0.16).

Lastly, we tested whether we observe a relationship between neural activity and LTM formation also for neuronal activity during the maintenance of stimuli from the non-preferred categories of cells (**Fig. 3g**) or for the visually evoked response of neurons when images were shown on the screen during encoding (**Fig. 3h**; encoding 1; 0 – 2s after picture onset; preferred trials only). However, none of the main effects nor interactions in either of the mixed-effects GLMs showed any significant relationship between FR and the factors tested (all p > 0.06).

### Activity during WM maintenance is linked to subsequent memory signal

During recognition memory tasks, a common observation in the MTL is that ‘memory selective’ (MS) cells differ in their response between novel and familiar images (Rutishauser et al. 2015). These cells are thought to represent a memory strength signal (Rutishauser et al. 2015), with stronger responses associated with stronger memories. We therefore next asked whether MS cells are present in the present experiment, and if so, whether there is a relationship between the activity of MS cells during LTM retrieval and that of category neurons during WM maintenance. Since we only observed a relationship between persistent activity of category neurons and successful LTM formation in the hippocampus, we restricted this analysis to the hippocampus.

We selected for MS neurons during all correct trials in the LTM recognition part (permutation-based t-test comparing response between correct familiar and novel images, that is, true positives with true negatives, 200-1200 ms after picture onset, p < 0.05). Out of the 351 recorded cells, n = 53 (15.10 %) were MS cells. Of these, 25 (47%) responded more to familiar than novel items, with the remaining responding more to novel than to familiar items. **Fig. 4a** shows an example neuron that increased its firing rate more for familiar than novel pictures (old > new). To assess whether MS cells, as expected, carry a memory strength signal, we next compared their response strength between trials retrieved with high and low confidence. We used receiver operating characteristics (ROC) analysis to do so (comparing old vs. new trials; see Methods). The response of MS neurons differed significantly more for high-as compared to low-confidence trials (**Fig. 4b**; t(40) = 2.89, p = 0.0039; 12 neurons excluded due to insufficient data in either one of the conditions, see Methods). This data shows that MS cells are present and signal memory strength in our experiment.

**Figure 4.**
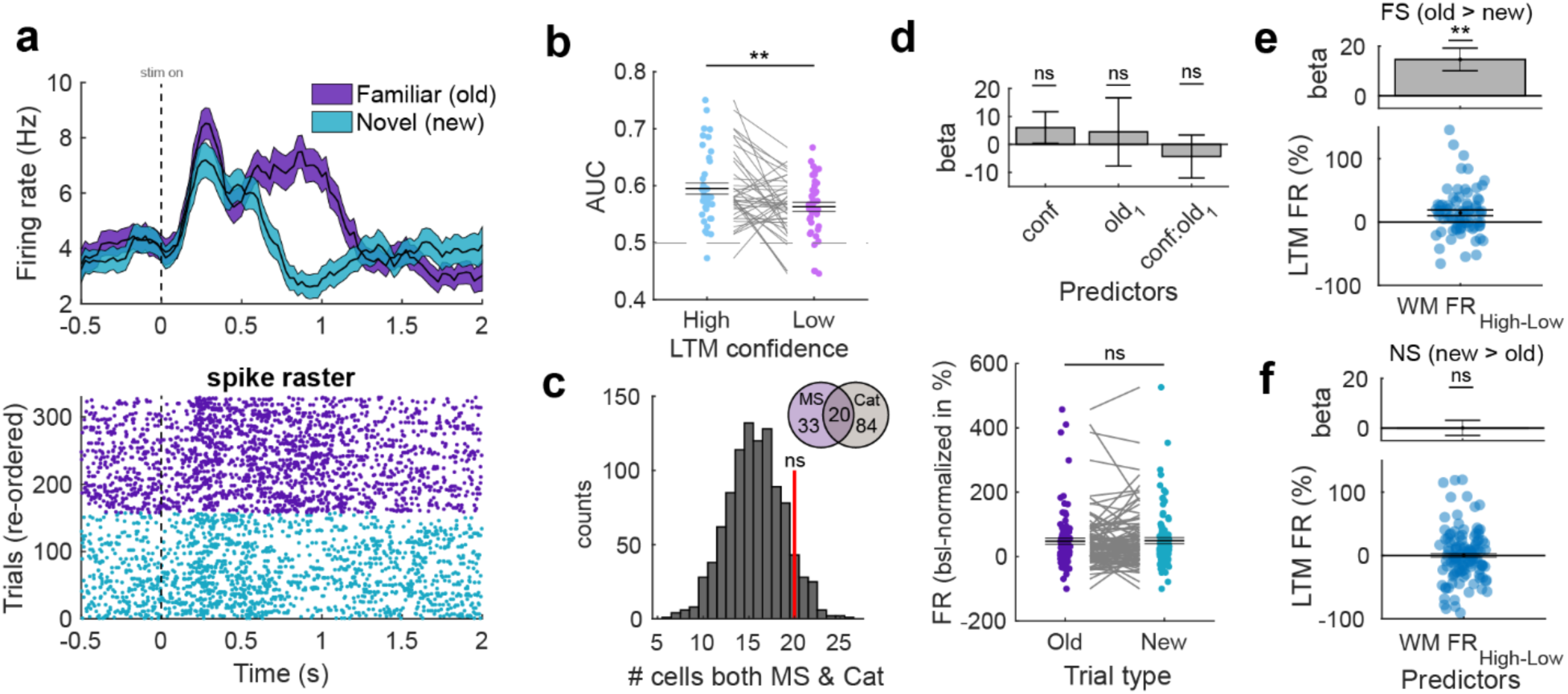
Relationship between memory-selective and category-selective neurons in the hippocampus. **(a)** Example of a memory-selective cell that had significantly higher FRs during correct old than new trials in the LTM recognition task. Colored areas in the PSTH plot (top) represent ± s.e.m. t=0 is stimulus onset. **(b)** ROC analysis comparing neuronal response of MS cells between new and old trials separately for low and high confidence. AUC was significantly higher for high than low confidence trials. Permutation-based t-test. Each dot is a neuron. **(c)** Null distribution of randomly selecting the same number of cells from the entire hippocampal population as category neurons for 1000 times and determining the overlap with MS cells. The overlap between category cells and MS cells (red bar) was not significantly higher than expected by chance. **(d)** Top: Mixed-effects GLM model testing for a relationship between FR of category neurons and *confidence* or *familiarity (“old_1-0_”)*. FR is estimated during picture presentation in the LTM task and only trials of the preferred category of a given cell are used. Bottom: Permutation-based t-test comparing FRs of category neurons during picture presentation between old and new items (preferred category only). Category neurons neither coded for familiarity nor novelty in the LTM task. **(e)** Response of familiarity-selective (FS) MS cells during familiar trials in the LTM task was stronger if maintenance activity for the same image was high during WM maintenance. Mixed-effects GLM. Top: Beta extracted from GLM result. Bottom: Distribution of FR differences between high and low maintenance trials. Each dot is a FS-category cell pair (n = 76). **(f)** Same as (e), but for novelty-selective (NS) MS cells (n = 122). The response of novelty-selective MS cells to familiar images did not differ significantly between whether the WM maintenance activity was low or high for an image. In **(b,d,e,f)** each dot is a neuron. In **(b,d**(bottom)) we used permutation-based t-tests. Center lines represent mean ± s.e.m. In **(d**(top)**,e,f)** error bars represent standard error of the coefficients. MS = memory-selective; Cat = category-selective; FS = familiarity-selective; NS = novelty-selective; ** p < 0.01, ns = not significant.

We next examined whether the overlap between the group of category neurons and MS neurons is significantly higher than chance, possibly hinting towards an involvement of category neurons also in LTM retrieval processes. Of the 53 MS neurons and the 104 category neurons, 20 neurons (37.8% of MS cells; 19.2 % of category neurons) were part of both groups. We determined whether this overlap is significantly higher that what would be expected by chance. We did so by randomly selecting the same number of neurons as we found for category neurons from the population of all hippocampal cells and determined how many neurons of those randomly selected neurons were also MS cells. This procedure was repeated for 1000 times to obtain a null distribution, used to determine the statistical significance of the originally observed overlap. This revealed that the overlap between category neurons and MS cells was not significantly higher than chance (**Fig. 4c**; p = 0.11), suggesting that category and MS cells were statistically independent populations of neurons (as expected (Rutishauser et al. 2015)).

To further examine whether, as a group, category neurons carried a memory signal during LTM retrieval, we computed a mixed-effects GLM with FRs of category neurons during all correct trials in the LTM recognition task (0.2 – 1.2 s after picture onset) in which the preferred category of a neuron was shown. We modelled *confidence* (3 levels, sure/unsure/guessing, continuous)*, familiarity* (2 levels, old/new, categorical), and their interaction as fixed effects. None of the effects revealed significant modulations of FRs (**Fig. 4d top**; confidence: beta = 6.05, p = 0.28; familiarity: beta = 4.49, p = 0.71; confidence x familiarity: beta = −4.32, p = 0.57). To further confirm this result, we directly compared FRs of category neurons between preferred old and new items and did not observe a significant difference (**Fig. 4d bottom**; t(94) = −0.28, p = 0.78).

Finally, we tested whether the response of MS cells in the recognition task to a given image was correlated with the activity of simultaneously recorded category cells during WM maintenance earlier in the same session while the same picture was held in mind. To do so, we examined all possible pairs of simultaneously recorded MS and category cells (n = 198). For each pair, we examined the trials during which images of the preferred category of the category cell were shown both in the WM and in the LTM task (i.e., preferred familiar trials; see Methods). We split the familiar trials in the LTM recognition task into two groups based on the level of activity of the category cell during WM maintenance for the same images (low vs. high maintenance activity, median split). We used a mixed-effects GLM to model FRs of the MS cells as a function of the fixed effect *Maintenance FR of category cells* (2 levels, high/low, categorical; based on simultaneously recorded FR of category neurons in the earlier WM task). We computed separate models for familiarity-selective (old > new) and novelty-selective (new > old) MS neurons as we hypothesized effects to be specific to neurons signaling familiarity (Rutishauser et al. 2015). This analysis revealed that FRs of familiarity-selective MS cells during LTM retrieval of items that were previously accompanied by high persistent activity in the WM task were higher than those previously accompanied by low persistent activity (**Fig. 4e**; Maintenance FR: beta = 14.63, p = 1.38 × 10^−3^). No significant relationship was observed for the activity of novelty-selective MS neurons (**Fig. 4f**; Maintenance FR: beta = 14.63, p = 1.38 × 10^−3^). This result suggests that the strength of category-selective WM maintenance activity is correlated with a neuronal measure of long-term memory strength (the activity of MS cells). This result is in addition to the correlation of WM maintenance activity and later behaviorally assessed long-term memory strength (previous paragraph).

## Discussion

Our results reveal that the activity of hippocampal category cells during WM maintenance was predictive of the success of LTM encoding. This relationship was specific for activity during the maintenance period and to trials in which the preferred category of category-selective cells was maintained in WM. In contrast, there was no significant correlation between neural activity of category cells during maintenance of stimuli from the non-preferred categories and LTM strength. There was no correlation between activity of the same category cells during the encoding period with later LTM memory strength, indicating specificity to activity during WM maintenance. Further, this effect was specific to the hippocampus as the activity of category neurons in the amygdala was not predictive of successful memory formation. Together, our findings reveal that the neural code used for maintaining items in WM is at least partially overlapping with the neural code that facilitates LTM encoding.

We also observed a relationship with a neuronal measure of LTM memory strength: the stronger the level of persistent activity for a given image, the larger was the response to that same image of memory-selective cells during the recognition memory test (**Fig. 4**). This reveals a direct neuronal-neuronal relationship between activity related to WM maintenance and LTM retrieval. Notably, this neuronal-neuronal relationship was only the case for the MS cells that increased their firing rate to familiar stimuli. In contrast, the MS cells that increased their firing rate to novel stimuli showed no significant correlation (for familiar stimuli). This result further supports the argument that what we observed is a signature of memory, because the activity of memory-selective cells scales with memory strength (and declared confidence) (Rutishauser et al. 2015).

Lesion studies indicate that the MTL is not necessary to perform simple WM tasks (Jeneson and Squire 2012), which has led to the long-standing idea of parallel memory systems, with the MTL not involved in WM. But if so, why is there persistent activity in the MTL during WM maintenance? One hypothesis is that the purpose of persistent activity is to engage mechanism used in encoding new memories in order be able to utilize synaptic plasticity to recover information in case it drops from the focus of attention (Kamiński and Rutishauser 2020). Under this framework, persistent activity would enhance the strength of items in LTM (Huang and Kandel 1994). This hypothesis is supported by more recent lesion studies, which show that subjects without a functional MTL do exhibit WM deficits in three situations: (1) in the presence of distractors (2), when memory load is high, or (3) when maintenance time is long (Jeneson and Squire 2012). In each of these scenarios, the probability that an item will drop out from the focus of attention is high and thus the network needs a mechanism for recovering this information. Here, we show that the extent of persistent activity in the hippocampus predicts whether items were encoded into LTM, thus revealing a specific example of a neural mechanism within the hippocampus that is engaged by both the WM and LTM system. We hypothesize that the role of persistent activity in the hippocampus is to augment the encoding of new information into LTM through repetition of the firing pattern throughout the maintenance period, thereby strengthening long-lasting long-term potentiation (Huang and Kandel 1994). This hypothesis is supported by theoretical work that indicates that the repetition provided by prolonged activity facilitates the modification of synapses (Jensen and Lisman 1996; Jensen et al. 1996).

The response properties of category cells in the amygdala and hippocampus were similar during WM processing but were remarkably different with respect to LTM encoding. In contrast to the hippocampus, activity of category cells from the amygdala did not predict LTM encoding success (**Fig. 3**). The relatively similar tuning properties of neurons in these two areas during encoding is not surprising in the context of prior work. For example, both brain areas contain concept cells (Quiroga et al. 2005, 2009) as well as MS cells (Rutishauser et al. 2008, 2015). Here, we now find that the relationship between short-term memory maintenance and its impact on later LTM is specific to the hippocampus. This is congruent with the fact that the hippocampus is particularly crucial for encoding new memories (Squire et al. 2004).

Our findings provide evidence for an interaction between WM and LTM where WM maintenance serves as a gating mechanism for LTM formation. In contrast, earlier research has shown that existing long-term memories can also be retrieved to and maintained in WM (Fukuda and Woodman 2017). In the current study, we used complex visual stimuli that are novel to the participants but could have pre-existing LTM associations for a specific person, animal, or object. It is thus possible that in our study such associations have been retrieved into WM to support the successful maintenance of these images. Since we didn’t make use of neutral images, such as fractals, without pre-existing associations or haven’t measured such associations for the current pictures on an individual basis, it is not possible for us to assess how LTM retrieval supported successful WM maintenance. We note, however, that even if this were the case, this would not explain our effect because regardless of whether WM maintenance engaged retrieval of existing images or not, encoding of a new memory was required to solve our task. We therefore interpret our findings as indicating that WM maintenance supported LTM formation. On a behavioral level, we observed that WM load had an influence on the successful formation of newly stored long-term memories, since images maintained in load 3 trials were less well remembered than images maintained in load 1 trials. Moreover, the strength of persistent activity during the maintenance period predicted successful encoding into LTM, but not the neural activity observed during encoding (see **Fig. 3**). These observations cannot be explained by LTM retrieval processes during the WM maintenance period. Nevertheless, we hypothesize that interactions between the systems likely go in both directions: persistent activity during WM maintenance predicts LTM formation, and, in turn, WM is supported by the retrieval of pre-existing LTM associations, presumably enhancing content-selective persistent activity during WM maintenance. However, future research is needed to shed more light on these interesting questions.

Our findings further suggest that the neural mechanisms of successful LTM formation overlap with those of WM maintenance. However, we emphasize that this does not mean that the two processes share exactly the same mechanisms such that the distinction between WM and LTM could eventually be discarded. Instead, our findings should be interpreted within a Hebbian view of two distinct memory systems: a short-term memory system that depends on reverberatory activity of cell assemblies and a long-term storage system that involves strengthening of synaptic connectivity between neurons (Nobre 2022). Our findings suggest that processes of LTM formation, which ultimately lead to successful LTM storage, become enhanced through interactions with persistent activity of WM-selective neurons. The exact mechanistic consequences of such interactions, however, remain the subject of future investigations. It also remains unclear whether other forms of WM maintenance, like activity-silent WM (Stokes 2015), interact with LTM formation in the same way as persistent neural activity. Activity-silent WM maintenance has mainly been observed for WM content outside the focus of attention (Rose et al. 2016; Wolff et al. 2017), which is an important difference from our study in which the focus of attention was not manipulated per item. Earlier research, however, indicated that attention to WM items enhances successful LTM formation (Hartshorne and Makovski 2019) which indicates that persistent activity plays a special role in interactions with LTM formation.

In conclusion, our study reveals that the activity of hippocampal category-selective cells during WM maintenance is predictive of LTM encoding success. This relationship is unique to the hippocampus and the WM maintenance period, with no similar predictive activity observed in the amygdala or during encoding periods. Our findings suggest that persistent activity of category cells in the hippocampus contributes to the encoding of declarative memories, reinforcing the role of Hebbian-type plasticity. They further show that WM- and LTM-specific neural populations interact on a local level with stronger persistent activity predicting stronger memory-related activity during retrieval. These results provide significant insights into the neural mechanisms involved in interactions between WM and LTM with single-cellular resolution.

## Acknowledgments

We thank the staff and physicians of the Epilepsy Monitoring Unit at Cedars-Sinai Medical Center for assistance, all patients and their families for their participation, Michael Kyzar, Nand Chandravadia, Andrea Schjetnan, and Ian Reucroft for spike sorting, data management, and data acquisition, Matthew Chen for behavioral prototyping, and Chrystal M. Reed and Jeffrey Chung for patient care. This work was supported by the National Institute of Health (U01NS117839 to U.R.), a Postdoctoral Fellowship by the German Academy of Sciences Leopoldina (to J.D.) and a Postdoctoral Award by the Center for Neural Science and Medicine at Cedars-Sinai (to J.D.).

## Author contributions

Conceptualization, J.D., J.K. and U.R.; Writing – Original Draft, J.D., J.K., and U.R..; Writing – Review & Editing, all authors. Investigation: J.D., J.K., Y.S.; Formal analysis: J.D. and U.R.; Methodology, J.D., J.K., and U.R.; Funding Acquisition, Resources, and Supervision, J.D., U.R. and A.N.M.; Performed surgery, A.N.M., T.A.V., and W.A.

## STAR Methods

### Resource availability

#### Lead Contact and Materials Availability

Further information and requests for resources should be directed to the Lead Contact, Ueli Rutishauser.

#### Data and Code Availability

Data will be made available publicly upon acceptance in the NWB format, similar to DANDI #673 (which contains only the WM part; we will add the LTM part). Example code will accompany the data release.

### Experimental Model and Study Participant Details

41 patients (48 sessions; 1 binary; 20 females; 20 males; age: 39.9 ± 12.9 years; **Table S1;** note of these, 38 sessions are also included in (Daume et al. 2024) and 10 were added here), undergoing invasive monitoring to assess treatment options for drug-resistant epilepsy, participated in the study. Two patients with task performance lower than 55% correct in either the WM or the LTM task were excluded from further analyses (see **Table S1**). All patients had Behnke-Fried hybrid electrodes (AdTech Inc.) implanted for intracranial seizure monitoring, gave their informed consent, and participated voluntarily. This study was part of an NIH Brain consortium between three institutions (Cedars-Sinai Medical Center, Toronto Western Hospital, and Johns Hopkins Hospital) and approved by the Institutional Review Board of the institution at which the patient was enrolled. Electrode localization was performed using a pre-operative MRI together with either MRI or CT post-operative images and Freesurfer as previously described (Minxha et al. 2020). Electrode positions are plotted on the CITI168 Atlas Brain (Tyszka and Pauli 2016) in MNI152 coordinates for the sole purpose of visualization (**Fig. 1b**). Coordinates appearing in white matter or outside of the target area is due to usage of a template brain. Electrodes that were localized outside of the target area in native space were excluded from analysis (4 out of a total of 149 recording sites).

### Method Details

#### Task

The study consisted of two separate tasks: a modified Sternberg WM task followed by a subsequent LTM recognition task. The WM task has been described elsewhere (Daume et al. 2024). It consisted of 140 trials and 280 novel pictures. In each trial, the onset of a fixation cross presented for 0.9 to 1.2 s (see **Fig. 1a top**) indicated the start of the trial. The fixation cross was followed by either one (load 1; 70 trials) or three (load 3; 70 trials) consecutively presented pictures, each presented for 2 s. A maintenance period of 2.55 to 2.85 s length followed the picture presentation, during which only the word “HOLD” was shown on the screen. Then, a probe picture was presented, which was either one of the pictures shown earlier in the same trial (match) or a picture already presented in one of the previous trials (non-match). The task was to indicate whether the probe picture matched on of the pictures shown earlier *in the same trial* or not. Note that the probe image shown was always one that had been shown before and thus familiar, with the answer ‘Yes’ if the image was shown in this particular trial and ‘No’ if it was shown in a previous trial. For trials where the correct answer was ‘No’ (i.e. the probe image was not shown during encoding in this trial), we used images that were presented in a previous trial to assure that all probe images were equally familiar, thereby preventing the use of novelty as a signal to answer the probe question. Probe images in the ‘No’ category were chosen from one of the categories for which no images were shown during encoding in a given trial. The probe picture was shown until patients provided their response via button press. All pictures shown during encoding were novel (i.e., the patient had never seen this particular image) and were drawn from five different visual categories: faces, animals, cars (or tools depending on the version), fruits, and landscapes. In load 3 trials, each image shown during encoding was from a different category.

After a brief delay (lasting 10 - 30min), patients completed a LTM recognition task. During this task, 400 images were shown one at a time. 200 of these images were new (not used in the WM task), whereas 200 were old (previously shown in the WM task). Each trial started with a fixation cross (**Fig. 1a, bottom**), followed by a single image for which the subject was asked to decide whether they had seen this image before (during the WM task) and to indicate the confidence in their response (sure, unsure, guessing). The image stayed on the screen until a response was given (no timeout). The ‘new’ images (foils) were chosen from the same 5 visual categories as the ‘old’ images. Note that due to this design, solving the recognition memory task required remembering the specific stimuli seen because the new images used were similar to the old images.

### Quantification and Statistical Analysis

#### Spike sorting

For each hybrid depth electrode, we recorded the broadband LFP signal between 0.1 and 8,000 Hz at a sampling rate of 32 kHz (ATLAS system, Neuralynx Inc.; Cedars-Sinai Medical Center and Toronto Western Hospital) or 30 kHz (Blackrock Neurotech Inc.; Johns Hopkins Hospital) from a total of eight microwires. All recordings were locally referenced within each recording site by using either one of the eight available micro channels or a dedicated reference channel with lower impedance provided in the bundle, especially when all channels contained recordings of neuronal spiking. We used the semiautomated template-matching algorithm OSort (version: 4.1) (Rutishauser et al. 2006) to detect and sort spikes from putative single neurons in each wire. Spikes were detected after bandpass filtering the raw signal in the 300 – 3,000 Hz band (see **Fig. S2** for single cell quality metrics). The two tasks (WM and LTM) were acquired in a single recording and all neurons were jointly sorted for both tasks. In total, we isolated 950 neurons across both areas of the MTL. Neurons with a firing rate lower than 0.1 Hz in either the WM or the LTM tasks were excluded from analysis (67 neurons (7.1%)). Analyses are based on 351 isolated neurons in the hippocampus and 532 in the amygdala (a total of 883 neurons across both areas).

#### Selection of neurons

To select for category neurons whose firing rate differed systematically between the picture categories during image presentation (encoding) in the WM task, we counted the number of spikes a given neuron fired in a window between 200 to 1,200 ms after picture onset across all trials in each category (all encoding periods and the probe period). We then then computed a 1×5 permutation-based ANOVA with visual category as the grouping variable. In addition, we computed a post-hoc right-sided permutation-based t-test between the category with maximum spike count and all other categories combined. We classified a given neuron as a category neuron if both tests were significant (both p < 0.05) (Daume et al. 2024). We refer to the category with the maximum average spike count as the preferred category of the cell. We note that category cells are selected only using spiking activity from the encoding period, leaving the firing rates during the maintenance period independent for later analyses.

In the recognition task, we selected for neurons that were memory-selective by comparing the number of spikes fired following image onset (window of 0.2-1.2 seconds after image onset) between correct familiar and novel trials using a permutation-based t-test (p<0.05, two-sided). Neurons with higher firing rates for familiar than novel items (old > new) were classified as familiarity-selective, the other way round (new > old) as novelty-selective.

#### Relating WM maintenance activity to LTM formation

We tested whether category-selective WM maintenance activity predicted LTM formation using a mixed-effects GLM across all category-selective cells. The analysis was performed on baseline-normalized firing rates during the maintenance period (0-2.5 s) of the WM task in trials in which the preferred category of a given neuron was held in WM. We only considered WM trials for which the probe question was answered correctly, and which contained an image that appeared in the subsequent LTM recognition test. We used LTM *accuracy* (2 levels, correct (remembered)/forgotten (wrong), categorical), LTM *confidence* (3 levels, high/medium/low, continuous), and *area* (2 levels, hippocampus / amygdala, categorical) as fixed effects and neuronID nested into patientID as random intercepts.

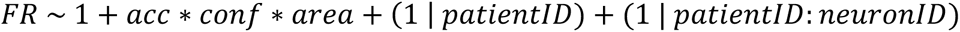

We hypothesized that firing rates during the WM maintenance period should be lower for pictures that were later forgotten (i.e., rated by mistake as “novel”) with high confidence (that is, “high-confidence wrong” trials) than those forgotten with low confidence (“low-confidence wrong” trials). The reason for this hypothesis is that for items that were forgotten with high confidence, there should be a weaker memory trace than for items for which patients were unsure whether they have seen the image before. We therefore labeled the confidence ratings for forgotten trials as high = 1, medium = 2, low = 3. This way the confidence labeling was consistent with our hypothesis. For remembered trials, in turn, we hypothesized that firing rates should be higher for high compared to low confidence trials, so we used the confidence labels high = 3, medium = 2, low = 1. To test whether category-selective activity predicted LTM formation during the encoding window, we based our GLM analysis on firing rates determined during the picture 1 window of preferred images (0-2 s).

#### Single-neuron AUC analysis

For MS neurons, we performed ROC analysis to assess how well the firing rate of individual cells distinguished between novel and familiar trials (Rutishauser et al. 2015). Spike counts between 200 and 1,200 ms after stimulus onset in the LTM recognition task were used for each neuron’s ROC analysis. We varied the detection threshold between the minimal and maximal spike count observed, linearly spaced in 25 steps. Only neurons with at least ten correct novel and familiar trials each were included. A separate ROC analysis was performed for high and low confidence trials. Only one of the two groups used for the ROC analysis was modified according to confidence while the other was kept constant. For familiarity-selective neurons, the fixed group was all true-negative trials (regardless of confidence) which was compared with high-confident true-positive and low-confident true-positive trials separately. For novelty-selective neurons, the fixed group was all true-positive trials which were compared with high-confident true-negative and low-confident true-negative trials separately.

#### Interaction between simultaneously recorded category- and memory-selective neurons

We determined whether MS neurons were more active during LTM retrieval of a familiar picture when the persistent activity of a simultaneously recorded category-selective neuron for that same picture was also high during the earlier WM maintenance period in the same session. To do so, we median-split the FRs of each category neuron during the maintenance period of correct trials that contained a preferred picture later tested in the LTM task into low and high FR trials (separately for load 1 and 3 trials to avoid a bias in FRs across loads). We then tested FRs of simultaneously recorded MS cells during LTM retrieval (determined during 0.2-1.2s after picture onset) between items that have been previously maintained with high vs low persistent activity. For that we used a mixed-effects GLM with *Maintenance FR* (2 levels, high vs low, categorical) as fixed effect and *neuronID* nested into *patientID* as random intercept.

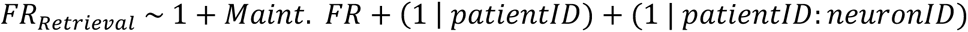

We performed this analysis separately for familiarity-selective (old > new) and novelty-selective (new > old) cells.

#### Statistics

For all statistical tests, we use (cluster-based) non-parametric permutation tests (statcond.m as implemented in EEGLab, using option ‘perm’, or ft_freqstatistics.m in FieldTrip), i.e., tests that do not make assumptions about the underlying distributions, or mixed-effects GLMs (fitglme.m in MATLAB) to assess statistical differences between conditions. Before each test, we removed neurons that differed ±3 SD from the mean across all neurons and all tested conditions. In the permutation-based tests, random permutations of condition labels were performed to estimate an underlying null distribution, which was then used to assess the statistical significance of the effect. All permutations statistics used 10,000 permutations, and t-tests were tested two-sided unless stated otherwise. The corresponding t estimates, which are computed based on a normal distribution, are provided as reference only. Cluster-based permutation statistics were performed as implemented in FieldTrip (Maris and Oostenveld 2007) with 10,000 permutations and an alpha level of 0.025 for each one-sided cluster. Lastly, error bars shown in figures reflect standard errors of the mean for permutation-based t-tests or standard errors of the coefficient for mixed-effect GLM results.

### Key Resources Table

See separate file.

