## Supplemental Information for "Persistent activity during working memory maintenance predicts long-term memory formation in the human hippocampus"

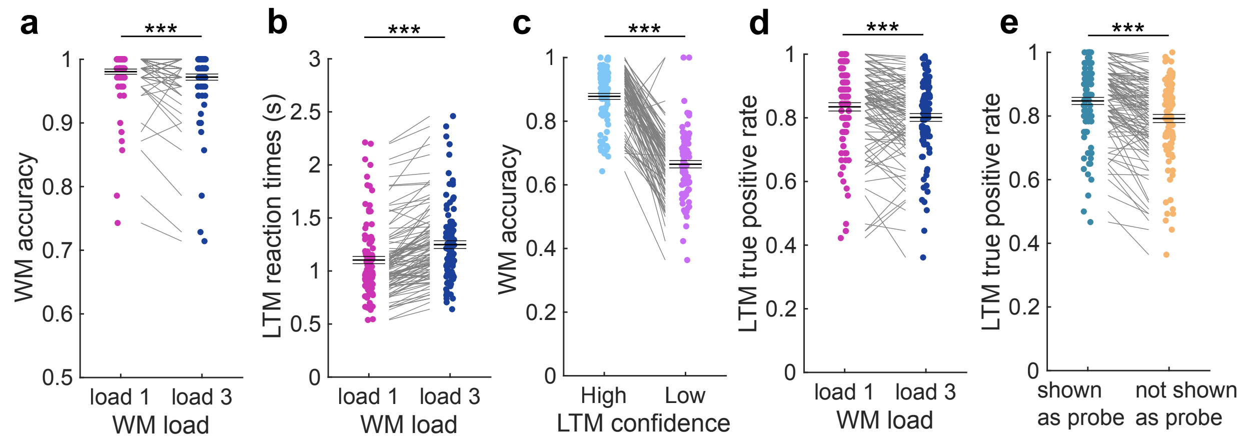


**Figure S1. Behavior results of 100 healthy control participants.** We conducted an online study collecting behavioral data from 100 healthy control participants (age: 32.9 ± 10.4 years; gender (self-reported): 54 females, 46 males). Participants were recruited on the online platform Prolific (www.prolific.co), gave informed consent, and were compensated monetarily for their participation ($4.50 base pay, bonus pay of $9.50 if performance above 75% in the WM task and 60% in the LTM task; total of $15 for approx. 1 hour experiment time; all but one participant received the bonus payment). We pre-selected participants based on age (18-80 years), location (USA), language (fluent English), and a minimum approval rate in previous studies (99%). The experiment was made accessible to participants through the online platform cognition.run (www.cognition.run) and programmed in JsPsych (version: 6.3.1). **(a,b)** In the WM task, participants responded **(a)** with higher accuracy ($t(99) = 3.54$; $p < 0.0001$) and **(b)** faster ($t(99) = -12.16$; $p < 0.0001$) in load 1 as compared to load 3 trials. **(c)** In the LTM task, participants remembered items better when they reported with high than low confidence. ($t(85) = 18.11$, $p < 0.0001$; 14 participants were not using the confidence rating and were therefore excluded from this analysis). **(d)** Pictures presented in load 1 trials in the WM task were better remembered in the subsequent LTM task than pictures presented in load 3 trials ($t(99) = 5.05$; $p < 0.0001$). **(e)** Pictures that were presented as probe images in the WM task were better remembered in the LTM task than pictures that were not used as probe images ($t(99) = 8.66$; $p < 0.0001$).

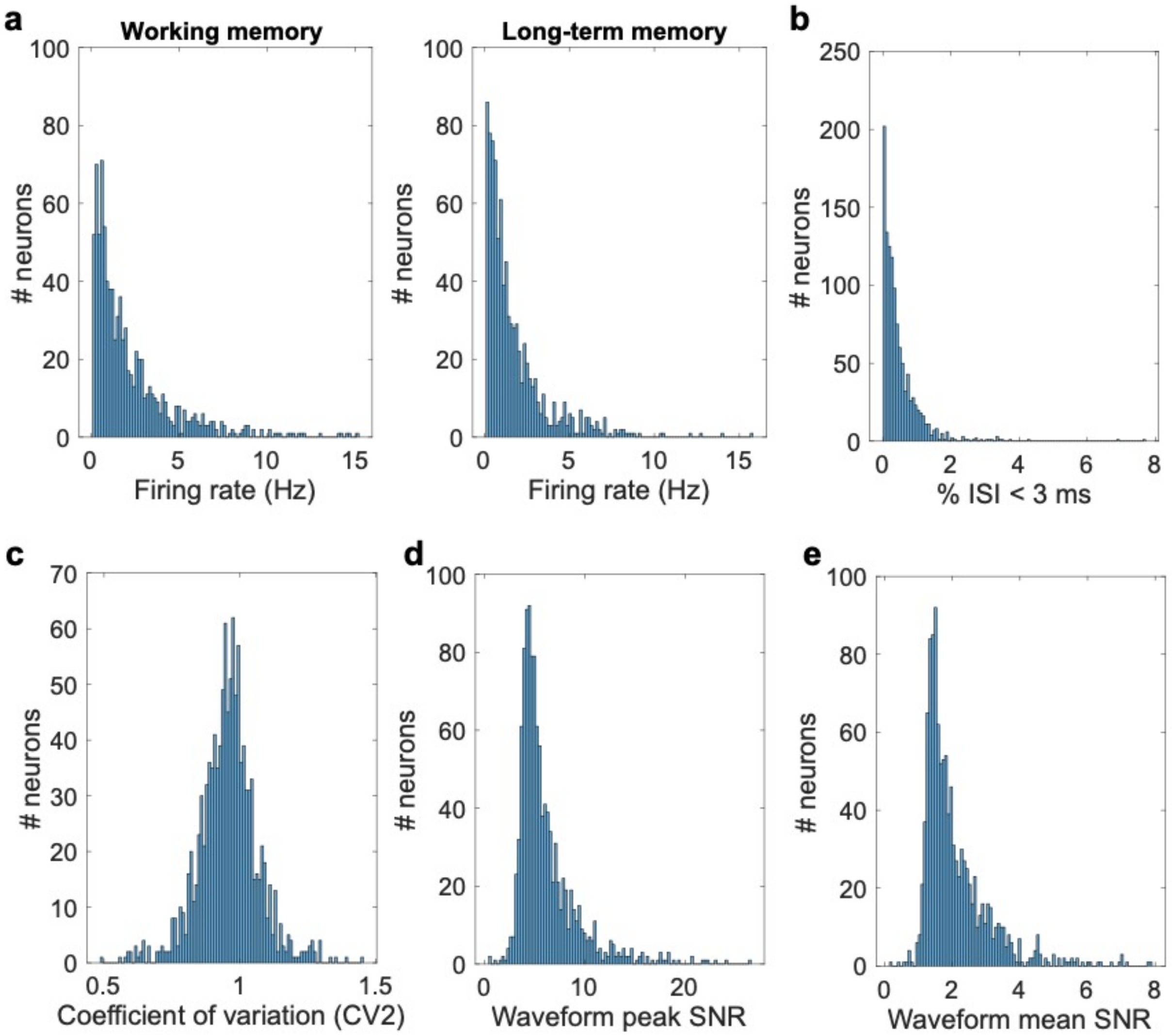


**Figure S2. Spike-sorting quality metrics for all identified putative single units. (a)** Average firing rate separately for the working memory (left) and the long-term memory part (right). **(b)** Proportion of inter-spike intervals (ISI) below 3 ms. **(c)** Coefficient-of-variation. **(d)** Signal-to-noise ratio (SNR) for the peak of the mean waveform across all spikes as compared to the standard deviation of the background noise. **(e)** Mean SNR of the waveform.

| Session | Gender | Age | Acc (%) (WM / LTM) | Seizure onset zone | Hippocampus | Amygdala |
|---|---|---|---|---|---|---|
| **P54cs_2** | f | 59 | 93.6 / 76.8 | Right mesial temporal | 3/0/7 | 5/2/17 |
| **P55cs** | f | 43 | 97.1 / 72.8 | Right mesial temporal | 0/0/1 | 7/3/21 |
| **P55cs_2** | - | - | 97.1 / 74.8 | - | 2/2/8 | 13/1/25 |
| **P57cs_2** | m | 46 | 91.4 / 67.5 | Left neocortical parietal | 1/0/4 | 14/5/24 |
| **P58cs** | f | 32 | 97.9 / 86.3 | Right frontal neocortical | 0/1/2 | 9/4/27 |
| **P61cs** | f | 52 | 87.1 / 68.8 | Left mesial temporal | 6/2/12 | 6/2/11 |
| **P61cs_2** | - | - | 97.9 / 72.3 | - | 0/0/2 | 7/3/8 |
| **P62cs** | f | 25 | 97.1 / 80.8 | Left mesial temporal | 0/2/3 | 5/0/19 |
| **P64cs** | f | 63 | 78.6 / 56.8 | Right lateral temporal neocortical | 0/0/0 | 3/0/17 |
| **P65cs** | f | 55 | 94.3 / 72 | Bilateral independent temporal | 5/0/9 | 7/3/18 |
| **P67cs** | f | 38 | 97.9 / 80.8 | Bilateral mesial temporal | 0/0/0 | 5/1/12 |
| **P68cs** | m | 54 | 92.1 / 58 | Bilateral mesial temporal | 3/3/16 | 1/0/17 |
| **P69cs** | f | 41 | 75 / 64.8 | Not localized | 3/2/8 | 4/0/7 |
| **P70cs** | f | 30 | 98.6 / 68.3 | Right temporal | 0/0/1 | 5/7/14 |
| **P70cs_2** | - | - | 96.4 / 73.9 | - | 0/0/1 | 0/0/5 |
| **P71cs** | m | 40 | 96.4 / 69 | Not localized | 0/0/0 | 0/0/3 |
| **P73cs** | f | 58 | 92.1 / 60.5 | Left mesial temporal | 0/1/5 | 7/1/17 |
| **P74cs** | m | 23 | 97.9 / 73 | Left neocortical temporal | 0/0/0 | 6/0/9 |
| **P76cs** | f | 24 | 99.3 / 78.5 | Not localized | 4/0/6 | 12/1/25 |
| **P76cs_2** | - | - | 100 / 90 | - | 0/0/0 | 0/0/4 |
| **P77cs** | f | 46 | 94.3 / 73 | Right auditory cortex | 2/2/4 | 12/12/41 |
| **P78cs** | f | 54 | 97.1 / 59.8 | Right anterior temporal | 0/1/13 | 0/1/14 |
| **P79cs** | f | 42 | 97.9 / 68.8 | Right anterior lateral temporal neocortex | 6/2/17 | 14/4/28 |
| **P79cs_2** | - | - | 97.1 / 80.5 | - | 4/0/13 | 9/3/19 |
| **P80cs** | o | 24 | 100 / 82.3 | Not localized | 4/7/13 | 20/9/49 |
| **P82cs** | m | 42 | 98.6 / 69.5 | Bitemporal | 11/2/17 | 12/1/24 |
| **P87cs*** | f | 26 | 74.3 / 54 | Not localized | 0/0/0 | 0/0/0 |
| **P88T** | m | 26 | 99.3 / 76.3 | Right mesial temporal | 3/1/22 | 0/0/6 |
| **P89T** | f | 45 | 97.1 / 57 | Right frontal | 7/3/22 | 11/0/18 |
| **P90T_2** | m | 20 | 91.4 / 57.8 | Occipital cortex | 0/0/11 | 0/0/2 |
| **P90T_3** | - | - | 90.7 / 55.8 | - | 0/1/13 | 0/0/0 |
| **P93T** | m | 23 | 83.6 / 62.7 | Left mesial temporal | 0/0/4 | 0/0/3 |
| **P96T** | f | 58 | 90 / 65 | Bilateral mesial temporal | 4/0/6 | 0/0/5 |
| **P101T** | f | 25 | 100 / 87 | Left neocortical temporal | 6/3/16 | 5/0/5 |

| | | | | | | |
|---|---|---|---|---|---|---|
| **P101T_2** | - | - | 100 / 90 | - | 5/3/8 | 5/3/6 |
| **P103T** | m | 49 | 96.4 / 61.5 | Right mesial temporal | 4/2/20 | 3/0/11 |
| **P106T** | m | 26 | 84.3 / 58.3 | Multifocal | 1/0/7 | 6/1/14 |
| **P109T** | m | 28 | 100 / 85.8 | Multifocal | 1/2/15 | 0/0/7 |
| **P110T** | m | 38 | 99.3 / 64.3 | Right fusiform cortex | 0/0/6 | 1/0/10 |
| **P113T_2** | m | 36 | 97.1 / 60.8 | Right mesial temporal | 2/2/11 | 1/1/7 |
| **P116T** | m | 28 | 97.9 / 65 | Left amygdala | 4/1/9 | 5/0/18 |
| **P1802jh** | m | 62 | 90.7 / 70.3 | Right mesial temporal | 5/1/12 | 0/0/0 |
| **P1809jh** | m | 45 | 75.7 / 57 | Left inferior + middle frontal gyrus | 0/1/6 | 0/0/0 |
| **P1811jh** | f | 27 | 76.4 / 61 | Bilateral mesial temporal | 1/0/10 | 0/0/0 |
| **P1814jh** | m | 48 | 90.7 / 76.8 | Left inferior orbitofrontal cortex and mesial temporal | 1/3/8 | 0/0/0 |
| **P1901jh*** | m | 55 | 45 / 74.8 | Left mesial temporal | 0/0/0 | 0/0/0 |
| **P1903jh** | m | 30 | 99.3 / 82 | Right mesial temporal | 0/0/2 | 0/0/0 |
| **P1912jh** | m | 49 | 70.7 / 60.5 | Left insula and left mesial temporal | 6/3/16 | 0/0/0 |

**Table S1. Patient demographics and neuron count per area.** For each area, the first number represents the category-selective neuron count, the second the memory-selective neuron count, and the third all recorded neurons. For sessions in which a patient performed lower than 55% in either of the two tasks, the neurons counts were set to zero (marked with *). These sessions were excluded from all analyses.
